## Supplementary Tables 1-3 for "Peptidoglycan recycling mediated by an ABC transporter in the plant pathogen *Agrobacterium tumefaciens*"

Running title: Peptidoglycan recycling mediated by an ABC transporter

Keywords: bacterial cell wall, peptidoglycan recycling, Agrobacterium tumefaciens, Rhizobiales, ABC transporter

### Table S1: Strains

| Bacterial Strain | Source |
|---|---|
| *Agrobacterium tumefaciens* C58 | - |
| *Eschericia coli* DH5α pNPTS139 | Courtesy of P. Brown |
| *Agrobacterium tumefaciens* C58 Δ*ampD* | This study |
| *Agrobacterium tumefaciens* C58 Δ*nagZ* | This study |
| *Eschericia coli* DH5α pSRKpTac | Courtesy of P. Brown |
| *Eschericia coli* DH5α pSRKpTac::*ldcA* | This study |
| *Agrobacterium tumefaciens* C58 pSRKpTac::*ldcA* | This study |
| *Agrobacterium tumefaciens* C58 Δ*ampD* pSRKpTac::*ldcA* | This study |
| *Eschericia coli* SM10 λPIR pSC189 | Chiang and Rubin, 2002 |
| *Agrobacterium tumefaciens* C58 Δ*yejABEF* | This study |
| *Agrobacterium tumefaciens* C58 Δ*yejABEF*Δ*ampD* | This study |
| *Eschericia coli* DH5α pSRKpTac::*ampG* | This study |
| *Agrobacterium tumefaciens* C58 Δ*yejABEF* pSRKpTac::*ampG* | This study |
| *Agrobacterium tumefaciens* C58 Δ*yejABEF*Δ*ampD* pSRKpTac::*ampG* | This study |
| *Agrobacterium tumefaciens* C58 Δ*ampC* | This study |
| *Agrobacterium tumefaciens* C58 Δ*yejABEF*Δ*ampC* | This study |
| *Agrobacterium tumefaciens* C58 Δ*yejA* | This study |
| *Agrobacterium tumefaciens* C58 Δ*yejB* | This study |
| *Agrobacterium tumefaciens* C58 Δ*yejE* | This study |
| *Agrobacterium tumefaciens* C58 Δ*yejF* | This study |
| *Agrobacterium tumefaciens* C58 Δ*yepA* | This study |

#### Table S2: Primers

| Primer | FCP Identifier | Sequence (5’ -> 3’) |
|---|---|---|
| ampD_UF | FCP3880 | GGCCAAGCTTCGCCCGTCTTTATAACCTCG |
| ampD_UR | FCP3881 | GAAATGTCGCTGGAAGGCAGGGGCAAAATCCGGCAGACAT |
| ampD_DF | FCP3882 | ATGTCTGCCGGATTTTGCCCCTGCCTTCCAGCGACATTT |
| ampD_DR | FCP3883 | GGCCGAATTCCACGAAGGTCTCCAGCGTCA |
| nagZ_UF | FCP3907 | GGCCAAGCTTGACATCAAGGCCGCCGAAAT |
| nagZ_UR | FCP3908 | CTTCAGCGCATTCATGGAGCATGAAACCCCAGGGCTGCTC |
| nagZ_DF | FCP3909 | GAGCAGCCCTGGGGTTTCATGCTCCATGAATGCGCTGAAG |
| nagZ_DR | FCP3910 | GGCCGAATTCAATACGCGACAGATCGACCT |
| yejABEF_UF | FCP4525 | AACCGGATCCCCAGCCGATCTTCTGCGCAT |
| yejABEF_UR | FCP4526 | ATTGAAGGCCGCTGCCAGCAGTTGGTGCCAATATCATCGGGG |
| yejABEF_DF | FCP4527 | CCCCGATGATATTGGCACCAACTGCTGGCAGCGGCCTTCAAT |
| yejABEF_DF | FCP4528 | AACCGTCGACTGCCTGATGGCTGCGGAATT |
| yejA_UR | FCP4529 | GATGAACATCTCGCCACGGTGGTGCCAATATCATCGGGG |
| yejA_DF | FCP4530 | CCCCGATGATATTGGCACCAACCGTGGCGAGATGTTCATC |
| yejA_DR | FCP4531 | AACCGTCGACATCGAAACCACCGGACTGGC |
| pSRK_ampG_F_NdeI | FCP4634 | CTTCATATGTCCAGTCAATATTTACGTATT |
| pSRK_ampG_R_HindIII_stop | FCP4783 | CTTAAGCTTTCACGTCAGATGCGTTTTTCGTA |
| ampC_U | FCP4630 | AACCGGATCCCCACGACCCCGAGAAACAGCAA |
| ampC_UR | FCP4631 | AGAGACGCGTTCCTCATTGGCCAAAGCGGCCAGTGCGATA |
| ampC_DF | FCP4632 | TATCGCACTGGCCGCTTTGGCCAATGAGGAACGCGTCTCT |
| ampC_DR | FCP4633 | AACCGTCGACGTAGAACCGAACATGGCCAC |
| pSRK_ldcA_F_NdeI | FCP4745 | GAACATATGTCTCTGTTTCACTTAATTGC |
| pSRK_ldcA_R_HindIII | FCP4746 | GAAAAGCTTTTACATTTTAAGAACAGGATG |
| yejB_UF | FCP4892 | GCGCCGGCCAGGCGCCAGAAGGCAACGAACAGGCGGATTT |
| yejB_UR | FCP4893 | AATCGATGCGCGGATCGATCCCGATCGTCGGGATCATCAGT |
| yejB_DF | FCP4894 | GGATCGATCCGCGCATCGAT |
| yejB_DR | FCP4895 | CGCGTTCGGCCGTGCTAGCGTTGCCAAGCGTACAGCCGGG |
| yejE_UF | FCP4273 | AACCGGATCCACCAGCTGAACTGGTGGCAGAAGAT |
| yejE_UR | FCP4274 | ATCTCACCCAGAGACGGCGATCTTACCAGCGTTTCGGTTCCGGC |
| yejE_DF | FCP4275 | GCCGGAACCGAAACGCTGGTAAGATCGCCGTCTCTGGGTGAGAT |
| yejE_DR | FCP4276 | AACCGTCGACATGGTGATGTCGTTGCCGCG |
| yejF_UF | FCP4888 | GCGCCGGCCAGGCGCCAGAAACCTGCTTTTGCAGCGTTTC |
| yejF_UR | FCP4889 | GTGTAATCCTGCTGCGGATTATCGACAGCGACGCTGGTTG |
| yejF_DF | FCP4890 | AATCCGCAGCAGGATTACAC |
| yejF_DR | FCP4891 | CGCGTTCGGCCGTGCTAGCGACACGGAGTGCTCGATCAGC |
| yepA_UF | FCP5281 | TCCTGCAGGATATCGTGGATCCAGGTCTTGAACCAGCCGAGC |
| yepA_UR | FCP5282 | AATGTAGTGGCCGGCCACGAGCATGGATATGCCGTGCAGCG |
| yepA_DF | FCP5283 | CTCGTGGCCGGCCACTACAT |
| yepA_DR | FCP5284 | AATACGACTCACTAGTGGGTCGACTTTCCTGAACGTCGCGGCCT |
| yejA_coexp_F | FCP6349 | GACGGCAAGCCGCGTGACATAT |
| yejA_coexp_R | FCP6350 | TTAACGCGGCCTTCCTTCACGG |
| yejB_coexp_F | FCP6351 | CGCTCACCCGTTTTCTTGAA |
| yejB_coexp_R | FCP6352 | GTCGAAGAATGATCCGCCGG |
| yejE_coexp_F | FCP6353 | GGATGAGATCAACGCCAATG |
| yejE_coexp_R | FCP6354 | AAATACCCCTGAATGGCGCC |
| yejF_coexp_F | FCP6355 | CTGCTGTTGCAGGTCGGCAT |
| yejF_coexp_R | FCP6356 | AACGATCTTGCCCTTGGTCA |

### Table S3: Identified Muropeptides

| Schematic | Composition | Ion $[M+H]^+$ | |
|---|---|---|---|
| | | Observed | Expected |
| 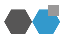 | GlcNAc-anhydroMurNAc | 479.1876 | 479.1872 |
| 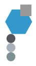 | anhydroMurNAc-L-Ala-D-Glu-m-DAP | 648.2722 | 648.2723 |
| 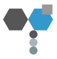 | GlcNAc-anhydroMurNAc-L-Ala-D-Glu-m-DAP | 851.3515 | 851.3517 |
| 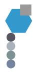 | anhydroMurNAc-L-Ala-D-Glu-m-DAP-D-Ala | 719.3102 | 719.3094 |
| 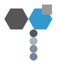 | GlcNAc-anhydroMurNAc-L-Ala-D-Glu-m-DAP-D-Ala | 922.3877 | 922.3888 |
| 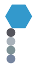 | MurNAc-L-Ala-D-Glu-m-DAP-D-Ala | 737.3199 | 737.3200 |
| 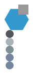 | anhydroMurNAc-L-Ala-D-Glu-m-DAP-D-Ala-D-Ala | 790.3455 | 790.3465 |
| 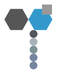 | GlcNAc-anhydroMurNAc-L-Ala-D-Glu-m-DAP-D-Ala-D-Ala | 993.4251 | 993.4259 |
| 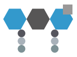 | MurNAc-(L-Ala-D-Glu-m-DAP)-GlcNAc-anhydroMurNAc-L-Ala-D-Glu-m-DAP | 1498.6155 | 1498.6166 |
| 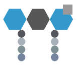 | MurNAc-(L-Ala-D-Glu-m-DAP-D-Ala)-GlcNAc-anhydroMurNAc-L-Ala-D-Glu-m-DAP-D-Ala | 1640.6883 | 1640.6909 |
| 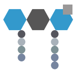 | MurNAc-(L-Ala-D-Glu-m-DAP-D-Ala)-GlcNAc-anhydroMurNAc-L-Ala-D-Glu-m-DAP-D-Ala-D-Ala | 1711.7255 | 1711.7280 |
| 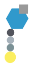 | anhydroMurNAc-L-Ala-D-Glu-m-DAP-D-Met | 779.3111 | 779.3128 |
| 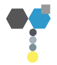 | GlcNAc-anhydroMurNAc-L-Ala-D-Glu-m-DAP-D-Met | 982.3940 | 982.3921 |
